# Intraspecific morphological variability of the invasive mosquito *Aedes koreicus* in Europe: genetic characterisation and population-level insights

**DOI:** 10.64898/2026.06.20.733498

**Authors:** Kornélia Kurucz, Safia Zeghbib, Ágota Ábrahám, Zsófia Tauber, Krisztián Bányai, Roger Eritja, Gábor Kemenesi

## Abstract

**Background:** The invasive mosquito *Aedes koreicus* has established populations in several European countries during the past decade, raising increasing public health concerns due to its potential role as a vector of pathogens. While species identification is primarily based on morphological characters, *Ae. koreicus* exhibits distinct morphological variants originating from mainland Korea and Jeju Island, which complicates surveillance and may lead to misidentification, particularly in regions where closely related species co-occur. To date, the genetic basis and population-level relevance of these morphological forms in Europe remain poorly understood.

**Methods:** We investigated the co-occurrence of two morphological forms of *Ae. koreicus* in Hungary, representing the first confirmed European location where both forms were detected sympatrically and even within the same breeding sites. Adult mosquitoes were morphologically characterised using diagnostic traits, and individuals representing both morphotypes were subjected to comprehensive genetic analyses. We sequenced multiple mitochondrial markers (COX1, COX2, COX3, ATP6, ND1, ND3) and the nuclear ITS2 region using Oxford Nanopore long-read sequencing. Phylogenetic reconstructions and haplotype network analyses were applied to assess genetic differentiation between morphotypes and to compare them with *Aedes japonicus* as a closely related reference species.

**Results:** Across all analysed mitochondrial and nuclear markers, no genetic differentiation was detected between specimens identified as the „mainland” or „Jeju-do” morphological forms of *Ae. koreicus*. Phylogenetic and haplotype network analyses consistently grouped individuals independently of their morphotype, indicating a shared genetic background at the population level. In contrast, *Ae. japonicus* formed a clearly distinct genetic lineage, confirming the robustness of the applied markers for interspecific discrimination.

**Conclusions:** Our results demonstrate that the observed morphological variability in *Ae. koreicus* populations are not underpinned by detectable genetic differentiation using commonly applied mitochondrial and genomic markers. These findings highlight the limitations of using morphology alone to infer population origin or structure and emphasise the need for heightened awareness of intraspecific variability in routine surveillance. Accurate morphological identification remains critical, particularly in citizen science–based monitoring programmes and AI-based automatization platforms, such as Mosquito Alert, to avoid confusion with morphologically similar invasive species. Further studies integrating genomic and ecological approaches are required to elucidate the mechanisms underlying morphological variability in this emerging vector species.

## Background

In recent decades, the global spread of invasive mosquito species has accelerated markedly, driven by increased international trade, human mobility, and environmental change. Several *Aedes* species have successfully colonised new regions outside their native ranges, establishing stable populations in temperate areas and posing growing challenges for vector surveillance and public health preparedness, particularly in urban and peri-urban environments (1). Among these emerging invaders, *Aedes koreicus* (Edwards, 1917) has attracted increasing attention since its first detection in Europe. Originally native to East Asia, this species has demonstrated a remarkable ability to tolerate cooler climatic conditions, facilitating its establishment in several European countries within a relatively short time span (2,3). As a container-breeding mosquito with a high degree of ecological plasticity, *Ae. koreicus* readily exploits artificial breeding sites, enabling its persistence in human-dominated landscapes and increasing the likelihood of contact with human populations (4). Although its vector competence has not been characterised as extensively as that of other invasive *Aedes* species, the increasing overlap of *Ae. koreicus* with human populations underscores the importance of accurate surveillance and risk assessment (5,6).

Accurate species identification is a cornerstone of mosquito surveillance, ecological studies, and epidemiological risk evaluation. In Europe, routine monitoring programmes rely predominantly on morphological identification, which remains indispensable for large-scale sampling efforts and for initiatives involving non-specialist contributors, including citizen science–based surveillance (7–9).

Morphological keys provide a rapid and practical tool for species identification, facilitating timely reporting and the implementation of appropriate control measures. However, the reliability of morphology-based identification depends on the stability and diagnostic value of the characters used. *Aedes koreicus* represents a particular challenge in this context due to its close resemblance to the related species *Aedes japonicus* (it was previously considered a subspecies or variety of *Ae. japonicus*), which even led to the risk of misidentification (10–12), and because of its documented morphological variability. In its native range, two forms slightly differing in morphology have been described, originating from the continental eastern Asian regions, including the Korean Peninsula, and from Jeju Island (Jeju-do), located off the southern coast of South Korea (Republic of Korea), respectively (12–14). These forms differ most noticeably in the number of white rings (stripes) on the hind legs: mosquitoes from the mainland have basal pale bands on hind tarsomeres I-IV., the tarsomer V. is usually entirely dark or show only a few pale scales (call it „mainland” morphotype), while the specimens from Jeju Island have basal pale bands (complete rings) also on hind tarsomer V. (call it „Jeju-do” morphotype) (13,14). Historically, these morphotypes have been considered as geographically separated populations within the native distribution of the species, and their diagnostic characters have been used to indicate geographic origin. On this basis, and according to DNA barcoding, the first population of *Ae. koreicus* that appeared in Europe (in Belgium, in 2008) was most probably introduced from Jeju Island (15). Since then, the emergence and/or dispersal of the species has been reported in several countries, in all cases linked to the population that appeared in Belgium, i.e. the „Jeju-do” form. Accordingly, this morphotype (characterised by basal pale bands on hind tarsomeres I-V.) was introduced in the guidelines for the surveillance of invasive mosquitoes in Europe by ECDC (European Centre for Disease Prevention and Control), serving as the diagnostic morphological characters of *Ae. koreicus* (16).

In 2016, however, another population of *Ae. koreicus* appeared in Germany, which clearly showed characteristics of the „mainland” morphotype (17). This was the first report of the morphological variant of *Ae. koreicus* from the mainland of Korea outside of its native range, also indicated an independent introduction of the species to Europe (17). Multiple introductions into Europe have since been supported by population genetic analyses (18); however, that study focused on the relationships among European populations and did not address the different morphological forms.

Notably, following the initial detection of *Ae. koreicus* in Hungary in 2016, represented solely by the „Jeju-do” morphotype in 2017-2018 (19,20), the „mainland” form was first recorded in 2019 in the country: larvae collected from used tyres as part of the local mosquito monitoring program in the city of Pécs (Baranya county), were reared to adulthood, and the emerged adults represented both morphological forms. Although the „mainland” morphotype was not detected in adult trapping surveys conducted that year (in 2019), and no field monitoring was performed in 2020, subsequent adult trapping from 2021 (via BG-Sentinel trap with CO_2_, Biogents AG, Germany) onwards confirmed the consistent presence of both „Jeju-do” and „mainland” morphotypes in the area. These findings represent the first documented coexistence of the two *Ae. koreicus* morphological forms outside their native range. Indeed, the origin of the Hungarian „mainland” form and its relationship with the „Jeju-do” population established in the area is unrevealed so far, requiring genomic analyses.

The primary aim of this study was to assess whether the distinct morphological forms of *Ae. koreicus* observed in Europe is supported by underlying genetic differentiation. Given the central role of morphology in routine mosquito surveillance and the documented variability of diagnostic characters in this species, clarifying the relationship between morphological traits and genetic background is essential for accurate species identification and for the reliable interpretation of monitoring data. To address this aim, we applied a combination of complementary genetic analyses that targeted both mitochondrial and nuclear markers. By integrating haplotype network, maximum likelihood and Bayesian phylogenetic analyses across multiple markers, our study provides a comprehensive and methodologically robust assessment of the genetic basis of morphological variability in *Ae. koreicus* populations in Europe.

## Methods

### Mosquito samples and morphological characterisation

Adult specimens of *Ae. koreicus* were collected from urban habitats in the city of Pécs, Hungary, using CO_2_-baited BG-Sentinel traps (Biogents AG, Germany), during the mosquito breeding season in 2021. The species and morphotype were confirmed based on the morphological characteristics of adult mosquitoes (15–17). Altogether 20 - 20 female individuals of both „mainland” and „Jeju-do” morphotypes were involved in the comparative study, which were described by observing (taking photos) the top and side views of the thorax and the hind legs, representing the key morphological characteristics (**Fig. 1**). Although the relatively low number of examined individuals and the single geographical origin may be considered limitations of the study, this sampling reflects the currently known distribution of the two morphotypes. To our knowledge, based on available international monitoring data and published records, this is currently the only known locality where both morphotypes have been found in sympatry; therefore, including additional samples or regions was not feasible.

**Fig. 1.**
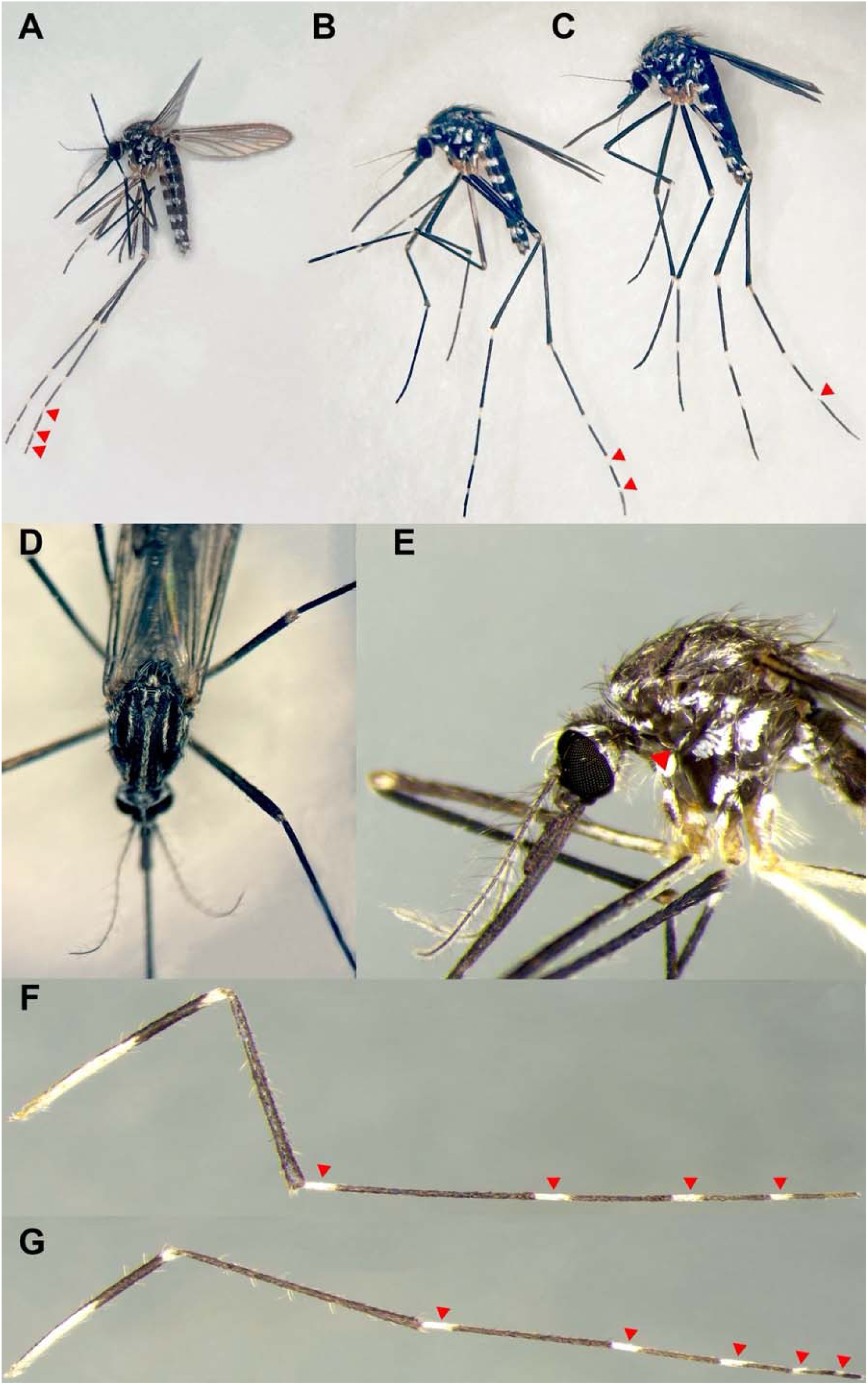
Key characteristics of different morphological forms of *Aedes koreicus* (**A:** *Ae. koreicus* „Jeju-do” morphotype, **B:** *Ae. koreicus* „mainland” morphotype, **C:** *Aedes japonicus*, **D - E:** *Ae. koreicus*’s thorax top and side views, representing both morphotypes, **F:** hind leg of *Ae. koreicus* „mainland” morphotype, **G:** hind leg of *Ae. koreicus* „Jeju-do” morphotype. Red triangles indicate the presence of basal pale scales (white rings) on hind leg tarsomers III – V.

### DNA extraction and sequencing

Total DNA from mosquitoes was extracted individually from the same specimens used for morphological analysis, using the NucleoSpin DNA Insect Kit (Macherey-Nagel GmbH, Germany), following the manufacturer’s instructions. Then, for further analyses, multiple mitochondrial DNA markers (COX1, COX2, COX3, ATP6, ND1 and ND3) were selected due to their widespread use in mosquito systematics, their sensitivity to demographic processes such as founder effects and population expansion, and their suitability for haplotype-based and phylogenetic analyses. In addition, the nuclear ITS2 region was included to provide an independent, biparentally inherited marker. Incorporating a nuclear locus enabled us to evaluate whether patterns observed in mitochondrial DNA were mirrored in the nuclear genome, thereby reducing the risk of overinterpreting maternally inherited signals and strengthening conclusions regarding population-level genetic structure.

The mitochondrial genes were amplified in two reactions using different primer pools (pool 1-2), designed to amplify ∼1500 basepair-long overlapping segments of *Aedes spp.* mitochondrion. Then, the ITS2 amplicons were generated separately, using the primers published by Manni et al. (21). In every case, we used Q5 Hot Start HF Polymerase (New England Biolabs, USA) in a 25 µl final volume. The details of designed primers, PCR reaction setups and conditions are available in the **Supporting information (S1-S3 Appendix**). Total volume of the mitochondrial pool 1-2 PCR reaction and 5 µl from the ITS2 PCR reaction were mixed before the products were cleaned up using AMPure XP Beads (Beckman Coulter, USA) with 80% ethanol for washing. The ends of the fragments were repaired using NEBNext Ultra II End Repair/dA-Tailing Module (New England Biolabs, USA). Barcodes from EXP-NBD196 (Oxford Nanopore Technologies, UK) kit were ligated with NEBNext Ultra II Ligation Module (New England Biolabs, USA) to the cleaned-up and end-repaired amplicons. Motor protein (AMX-F) from the LSK-110 kit (Oxford Nanopore Technologies, UK) was ligated to the barcoded fragments using NEBNext Quick Ligation Module (New England Biolabs, USA). For the quality check, Qubit Fluorometer was used with 1X dsDNA High sensitivity kit (Invitrogen, USA). The final library was loaded onto an R9.4.1 (Oxford Nanopore Technologies, UK) flow cell and sequencing run on the MK1C device for 72 hours. We successfully retrieved the targeted 7 markers from all the subject samples. The individual coverage of gene/sample was greater than 100x in every case.

A super-accurate basecalling model was used to generate FastQ files from raw Fast5 data with Guppy Basecaller (v6.4.6). The overall quality was measured with NanoPlot, and the reads were quality filtered using NanoFilt (22). The mean quality score (Q Score) reached 12.6, and unique reads above Q Score 11 were accepted. Demultiplexing was performed using Guppy Barcoder (v6.4.6) with default parameters and the requirement of barcode both ends and trimming the barcodes. The whole data under the same barcode was merged and assembled to distinct amplicon sequences using the Amplicon_sorter (23). Amplicon sequences were accepted if there was a build-up from more than 100 individual reads. The sorted amplicons were polished with the quality-filtered reads using Medaka (1.7.2). The polished mitochondrial (COX1, COX2, COX3, ATP6, ND1 and ND3) and genomic (ITS2) consensus sequences were mapped to an *Ae. koreicus* complete mitochondrial genome (Accession number: MZ460582.1) and available ITS2 sequences (Accession numbers: HG763830.1, FJ403046.1, PP835557.1), respectively, with Minimap2 (v. 2.23) (24). The consensus sequences were extracted using Geneious Prime (v.2023.0.4). The whole processing and genome assembly were conducted under the Ubuntu 22 operating system.

### Haplotype network, Maximum Likelihood and Bayesian Phylogenetic analyses

For the gene-by-gene comparison, sequences from the same region were aligned using the MUSCLE algorithm (25). As part of the haplotype network reconstruction, aligned sequences were first analysed in DnaSP v6 to identify unique haplotypes (26). The output haplotype data were exported in NEXUS format and manually edited following the POPART v1.7 (Population Analysis with Reticulate Trees) template to include morphotype (Mainland Korea or Jeju-do) as a character trait. Subsequently, the modified NEXUS file was imported into POPART v1.7, where the Median-Joining (MJ) algorithm, with epsilon set to 0, was used to infer relationships among haplotypes (27), This analysis was performed separately for each marker to visualise fine-scale genetic relationships and assess haplotype sharing between morphological forms, providing an intuitive framework for detecting potential population structure or lineage segregation.

In addition, to evaluate genetic relationships within a tree-based context and to investigate the presence of morphotype-associated clustering across multiple markers, phylogenetic analyses based on maximum likelihood and Bayesian inference were employed. The use of both approaches allowed us to assess the robustness of inferred patterns under different model assumptions and to identify well-supported versus poorly resolved relationships.

Bayesian phylogenetic reconstruction was performed with PhyloBayes-MPI using the Bayesian Markov chain Monte Carlo tree-sampling method, applying the CAT+GTR model uniformly across all markers (28,29). Supports at nodes were accepted if the posterior probabilities exceeded 0.95. Maximum difference was under 0.1 except in the case of COX1 (∼0.13) and ND1(∼0.11). For the visualisation, the iTOL webserver was used (30).

## Results and Discussion

**The haplotype network analyses** based on individual mitochondrial markers (COX1, COX2, COX3, ATP6, ND1 and ND3) consistently revealed extensive haplotype sharing between *Ae. koreicus* specimens assigned to the „mainland” and „Jeju-do” morphological forms (**Fig. 2, A-F**). Across all mitochondrial loci, haplotypes representing the two morphotypes were mixed within the same network structures, with no haplotype or haplotype cluster uniquely associated with either morphological form. Several haplotypes were shared by individuals of both morphotypes, including central haplotypes that accounted for a substantial proportion of the analysed specimens.

**Fig. 2.**
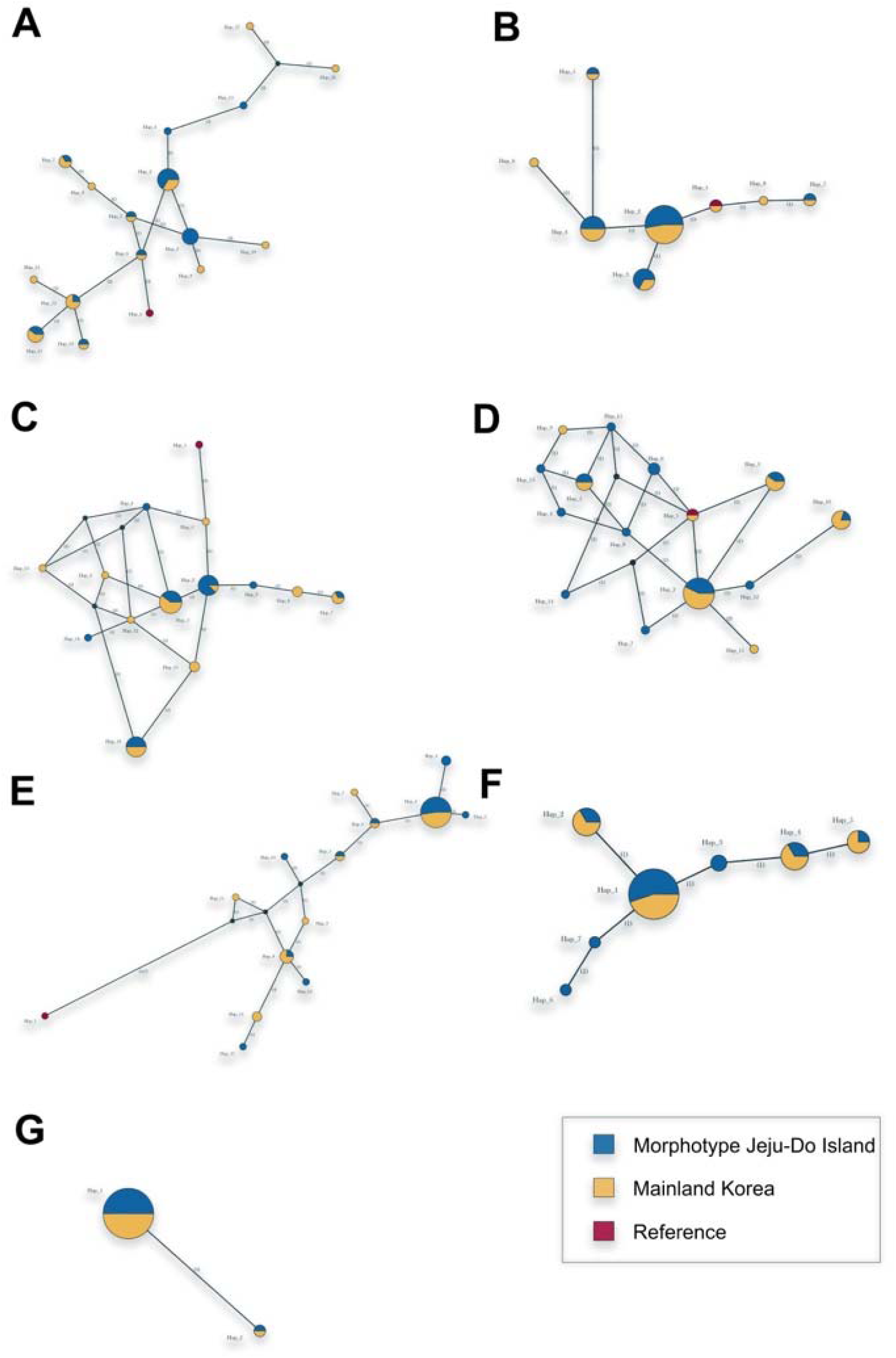
Haplotype networks of *Aedes koreicus* based on different genetic markers (**A:** COX1, **B:** COX2, **C:** COX3, **D:** ATP6, **E:** ND1, **F:** ND3, **G:** ITS2) of the Hungarian population. Different colours represent different morphological forms of *Ae. koreicus* (blue – „Jeju-do” morphotype, yellow – „mainland” morphotype, red – *Aedes japonicus* as the reference species).

The topology of the mitochondrial haplotype networks was characterised by low overall genetic divergence, with haplotypes separated by few mutational steps. In several markers, star-like or weakly branched network configurations were observed, which reflects limited mitochondrial diversity within the analysed dataset. Importantly, individuals representing different morphological forms frequently shared identical or closely related haplotypes, indicating a common mitochondrial genetic background across morphologically distinct specimens. In contrast, *Ae. japonicus*, included as a closely related reference species, consistently formed a clearly separated haplotype cluster, confirming the discriminatory power of the mitochondrial markers at the interspecies level.

At this spatial scale, reduced mitochondrial diversity and extensive haplotype sharing are consistent with a recent introduction and local population expansion, in which insufficient time has elapsed for mitochondrial differentiation to accumulate. Additionally, mitochondrial analyses highlighted the limited utility of the used markers in determining the specimens’ origin and genetic discrimination between the two morphotypes, as no clear association was observed between haplotypes and morphotype. The presence of minor mitochondrial variation within the Hungarian population may reflect standing variation introduced during one or more introduction events, subsequent accumulation of mutations after establishment, or repeated introductions from genetically similar source populations; however, the current dataset does not allow these scenarios to be distinguished.

The analysis based on the nuclear ITS2 marker showed results consistent with the mitochondrial data, since ITS2 sequences showed no differentiation between specimens assigned to different morphological forms of *Ae. koreicus*, with identical or near-identical ITS2 variants detected across both morphotypes (**Fig. 2, G**). No ITS2 sequence variant was exclusively associated with either the „mainland” or „Jeju-do” morphological form.

Supporting the patterns observed in the mitochondrial analyses, *Ae. japonicus* formed a distinct and clearly separated ITS2 lineage, further validating the suitability of this marker for interspecific discrimination.

**Maximum likelihood phylogenetic analyses** based on both the mitochondrial markers (COX1, COX2, COX3, ATP6, ND1 and ND3) and the nuclear ITS2 region revealed highly similar patterns across all loci (**Fig. 3**). In each analysis, *Ae. koreicus* specimens formed a single lineage, with no phylogenetic separation corresponding to the „mainland” or „Jeju-do” morphological forms. Individuals representing different morphotypes were consistently distributed throughout the phylogenetic trees, and no morphotype-specific clades or marker-dependent clustering patterns were observed.

**Fig. 3.**
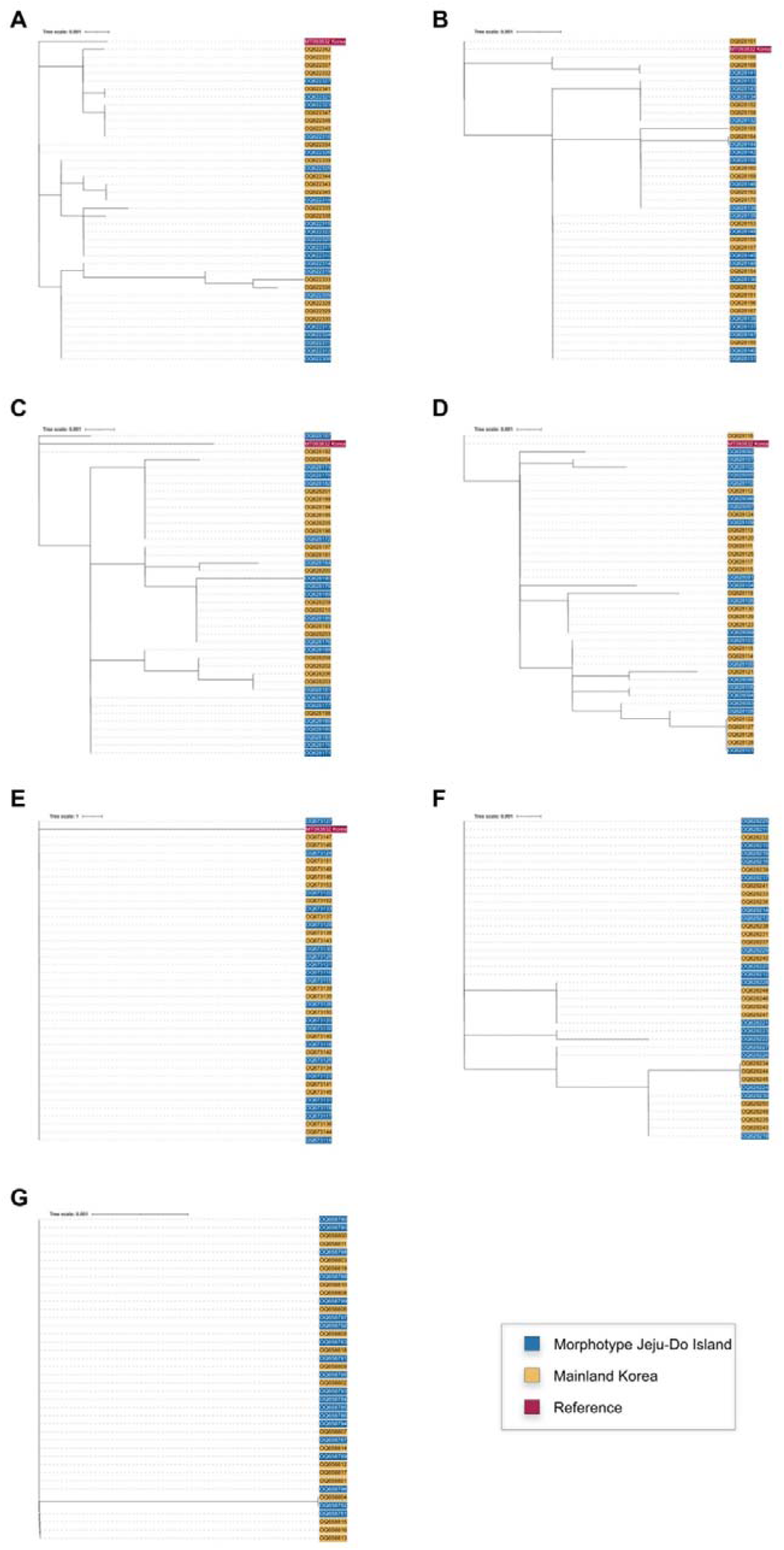
Maximum likelihood phylogenetic analyses of *Aedes koreicus* based on different genetic markers (**A:** COX1, **B:** COX2, **C:** COX3, **D:** ATP6, **E:** ND1, **F:** ND3, **G:** ITS2) of the Hungarian population. Different colours represent different morphological forms of *Ae. koreicus* (blue – „Jeju-do” morphotype, yellow – „mainland” morphotype, red – *Aedes japonicus* as the reference species).

Across all mitochondrial markers, branch lengths within the *Ae. koreicus* clade were short, indicating low levels of genetic divergence among the analysed specimens. This pattern closely mirrors the results obtained from haplotype network analyses and reflects limited genetic diversity within the dataset. Importantly, specimens representing different morphological forms frequently clustered together, further supporting the absence of phylogenetic structure associated with morphological variability (**Fig. 3, A-F**).

The phylogenetic tree based on the nuclear ITS2 marker showed a similarly undifferentiated pattern. All *Ae. koreicus* ITS2 sequences clustered into a single group, with no separation between morphological forms. The concordance between mitochondrial and nuclear markers strengthens the inference that the observed morphological variability is not linked to underlying genetic differentiation (**Fig. 3, G**). While mitochondrial DNA is maternally inherited and particularly sensitive to demographic processes such as founder effects, the nuclear ITS2 region provides complementary biparental information (31). The lack of differentiation across both marker types, therefore, argues against cryptic population structure associated with morphotype identity. In this case also, the reference taxon *Ae. japonicus* consistently formed clearly separated and well-supported lineages in all phylogenetic analyses.

**Bayesian phylogenetic analyses** performed on the same set of mitochondrial markers (COX1, COX2, COX3, ATP6, ND1 and ND3) and the nuclear ITS2 region produced tree topologies consistent with those obtained from maximum likelihood analyses. Within the *Ae. koreicus* clade, individuals representing different morphotypes were distributed throughout the Bayesian trees, and no morphotype-specific clusters were observed. Indeed, the reference sequences of *Ae. japonicus* formed clearly separated and well-supported lineages, indicating that the Bayesian framework was also able to resolve interspecific relationships where genetic differentiation was present (**Fig. 4**). The absence of morphotype-associated structure was consistent across mitochondrial and nuclear datasets, further supporting a shared genetic background among morphologically distinct individuals. Considering the spatial and temporal scale of our samples, limited genetic variation and extensive allele sharing are typical, and Bayesian inference is unlikely to recover well-supported internal structure unless strong differentiation is present. Hence, the Bayesian phylogenetic analyses reinforce the conclusions drawn from maximum likelihood and haplotype network approaches, providing additional support for the absence of genetic differentiation between the morphological forms of *Ae. koreicus* detected in Hungary.

**Fig. 4.**
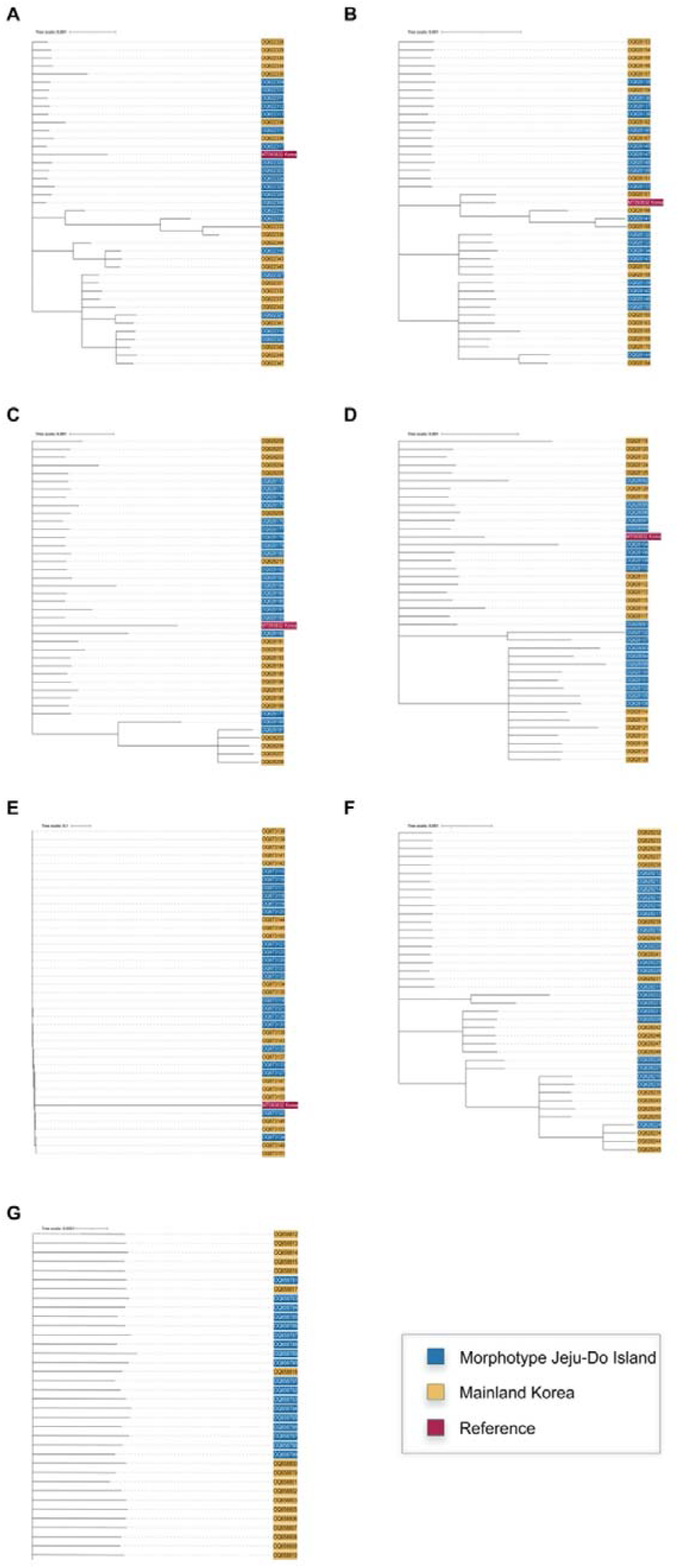
Bayesian phylogenetic analyses of *Aedes koreicus* based on different genetic markers **(A:** COX1, **B:** COX2, **C:** COX3, **D:** ATP6, **E:** ND1, **F:** ND3, **G:** ITS2) of the Hungarian population. Different colours represent different morphological forms of *Ae. koreicus* (blue – „Jeju-do” morphotype, yellow – „mainland” morphotype, red – *Aedes japonicus* as the reference species).

Although concatenation of mitochondrial markers could increase the number of informative sites, these loci represent a single maternally inherited linkage unit and therefore do not provide independent evidence of recombination or biparental population structure. For this reason, the separate-marker approach used here provides a conservative assessment of whether morphotype-associated genetic structure is consistently detectable across loci. The consistent results of Bayesian, maximum likelihood and haplotype network-based analyses across several maternally inherited mitochondrial markers and the biparentally inherited nuclear ITS2 region indicate that the morphological forms of *Ae. koreicus* described from the native range do not correspond to genetically distinct lineages within the examined Hungarian population. Thus, in this dataset, morphotype identity does not appear to reflect underlying genetic subdivision at the population level. Also, based on the available data, we cannot support the hypothesis that the co-occurrence of distinct morphological forms in Hungary reflects independent introduction events from different source regions, such as mainland Korea and Jeju Island.

Instead, an alternative explanation for the observed morphological variability may be phenotypic plasticity of the invasive population, retained ancestral variation, or environmentally influenced expression of phenotypic traits. In particular, the presence or absence of the additional pale band on the hind tarsomere may represent a developmentally plastic trait influenced by larval environmental conditions rather than a stable marker of geographic ancestry, although this hypothesis requires confirmation under controlled rearing conditions. Such patterns (similar morphological traits arise independently under comparable environmental conditions) have been widely documented in invasive insects and do not necessarily imply shared geographic origin or deep genetic divergence (32,33). Broader sampling from both the invaded European range and the native range, including reference sequences from mainland Korea and Jeju Island, would be necessary to test whether Hungarian haplotypes can be linked to particular source regions and to reconstruct the introduction history of *Ae. koreicus* with greater confidence. Indeed, our findings require further interpretation, as the variability within *Ae. koreicus* is more likely to be ecological rather than genetic in origin.

From a surveillance perspective, these results highlight the importance of interpreting morphological variation and the need for a refined knowledge of the diagnostic characteristics used in routine identification. Morphology-based identification remains crucial in surveillance programs, provided that diagnostic keys adequately reflect intraspecific variability. We therefore recommend that species identification guidelines, particularly those used in European surveillance frameworks (e.g. ECDC (16)), be updated to include both known morphological forms of *Ae. koreicus*.

Increased attention should be paid to morphological determination in future surveillance activities, especially in citizen science approaches, where identification of mosquito species is based on photographs (8). Given the close morphological similarity between *Ae. koreicus* and *Ae. japonicus*, along with their overlapping breeding habitat preferences (34), we further recommend retrospective re-examination of archived *Ae. japonicus* material from regions at risk of *Ae. koreicus* establishment. Such efforts would improve early detection, reduce the risk of misidentification, and ultimately enhance the accuracy and reliability of invasive mosquito surveillance across Europe. Until the heritability and environmental sensitivity of the diagnostic tarsal banding pattern are tested experimentally, morphotype identity should be treated as a potentially variable phenotypic character rather than as a reliable proxy for geographic origin.

At the same time, the demonstrated co-occurrence of distinct morphological forms of *Ae. koreicus* within a single European population, without detectable genetic differentiation, indicates that morphological variability alone should not be used to infer population origin or invasion history. This emphasises the practical value of integrating genetic data into mosquito monitoring programmes, particularly in regions where multiple invasive *Aedes* species co-occur.

Beyond its relevance for surveillance programs and morphological species identification, the recognition of morphotype-level variation in *Ae. koreicus* may also have implications within a One Health framework. The vectorial capacity of *Ae. koreicus* remains poorly understood, and phenotypic differentiation could add a further layer of complexity to the assessment of its epidemiological relevance. Although currently speculative, potential differences in vector competence, host-seeking behaviour, ecological tolerance, or seasonal dynamics among morphotypes should not be overlooked. Future studies integrating morphological, genetic, ecological, and vector competence data would therefore be valuable to clarify whether morphotype differentiation has any functional significance for public, animal, or environmental health risk assessment.

## Conclusions

This study provides, to our knowledge, the first integrated assessment of morphological variability and its genetic background in relation to the most common genetic markers in an invasive European population of *Ae. koreicus*. By demonstrating the co-occurrence of distinct morphological forms within a single Hungarian population without detectable mitochondrial or nuclear genetic differentiation at the population level, our results challenge the assumption that these forms necessarily reflect separate evolutionary lineages or independent invasion events.

Our findings indicate that morphological variation in *Ae. koreicus* is not associated with the population genetic structure detectable using the markers analysed here, highlighting the limits of morphology-based inference for reconstructing invasion history. At the same time, they emphasise that intraspecific morphological variability is a biologically meaningful feature of invasive populations and should be explicitly considered in surveillance frameworks.

In conclusion, this work underscores the need for an integrated approach to invasive mosquito monitoring that combines refined morphological knowledge with genetic data. Such an approach is essential to ensure accurate species identification, avoid misinterpretation of phenotypic variation, and strengthen early detection and surveillance of invasive *Aedes* species across Europe.

## Supporting information

Supporting information Apendix S1

## Acknowledgements

We would like to thank Gábor Endre Tóth, Zsaklin Varga and Rebeka Csiba for their assistance in the preparation of samples. We would acknowledge the Biological and Sportbiological Doctoral School of the University of Pécs, Hungary, for providing research opportunities for ÁÁ during her PhD studies.

## Availability of data and materials

All data used to generate conclusions are represented in figures and Supplementary information.

## Authors’ contributions

KK: conceptualisation, supervision of the investigations, collection of mosquito samples and species identification, laboratory processes, writing the original draft. ÁÁ: laboratory processes, sequencing. ZsT and SZ: bioinformatics, data analyses, visualisation. KB and RE: review and edit the manuscript. GK: conceptualisation and review of the manuscript.

## Ethics approval and consent to participate

Not applicable.

## Consent for publication

All the authors consent to the publication of the manuscript.

