## Supporting information Apendix S1 for "Intraspecific morphological variability of the invasive mosquito *Aedes koreicus* in Europe: genetic characterisation and population-level insights"

**S1 Appendix.** Primers used for mitochondrial genes (COX1, COX2, COX3, ATP6, ND1, and ND3) and nuclear ITS2 gene amplification.

| Primer Name | Primer Sequence 5'->3' | Organelle | Reaction Pool | Reference |
|---|---|---|---|---|
| aedes_mito_1500_2_LEFT | AGCCACCCTGGTATATTTATTGGG | mitochondrial | 1 | this study |
| aedes_mito_1500_2_RIGHT | GTCCTAAATTTGCTCATGTTGCCA | mitochondrial | 1 | this study |
| aedes_mito_1500_3_LEFT | ACACTTCCTCCTGCAGAACATAC | mitochondrial | 2 | this study |
| aedes_mito_1500_3_RIGHT | TGTTGTGTGTGACAAATTCATCCA | mitochondrial | 2 | this study |
| aedes_mito_1500_4_LEFT | GGCCCTAATGGACATAATGGAAGA | mitochondrial | 1 | this study |
| aedes_mito_1500_4_RIGHT | GGGTCGAATCCACATTCAAATGG | mitochondrial | 1 | this study |
| aedes_mito_1500_5_LEFT | GCAGCTGCTTGATATTGACATTTTG | mitochondrial | 2 | this study |
| aedes_mito_1500_5_RIGHT | CAGGGTTAACTGTATGTTATTCTTTTCGA | mitochondrial | 2 | this study |
| aedes_mito_1500_10_LEFT | CCAAATAAATTAGGAGGGGTAATTGCA | mitochondrial | 1 | this study |
| aedes_mito_1500_10_RIGHT | TCTGAGTTCAAACCGGCGTAAG | mitochondrial | 1 | this study |
| aedes_mito_1500_11_LEFT | AGCAAATCCCCCTCTTCTATATTCT | mitochondrial | 2 | this study |
| aedes_mito_1500_11_RIGHT | TGAAAGGTTTAAATAAGGAATTCGGCA | mitochondrial | 2 | this study |
| aedes_mito_1500_12_LEFT | TGCTACCTTCGCACAGTCAAAA | mitochondrial | 1 | this study |
| aedes_mito_1500_12_RIGHT | CCAGCTACCGCGGTTATACAAA | mitochondrial | 1 | this study |
| Ae_alb_ITS2_F | ATCACTCGGCTCGTGGATCG | nuclear | ITS2 | Manni et al. 2015 doi.org/10.1186/s13071-015-0794-5 |
| Ae_alb_ITS2_R | ATGCTTAAATTTAGGGGGTAGTCAC | nuclear | ITS2 | Manni et al. 2015 doi.org/10.1186/s13071-015-0794-5 |

**S2 Appendix.** Reaction setup and conditions for pools 1 and 2 (amplifying mitochondrial genes COX1, COX2, COX3, ATP6, ND1, and ND3)

| **Component** | **Amount for 25 µl reaction** |
|---|---|
| 5X Q5 Reaction Buffer | 5 µl |
| 10 mM dNTPs | 0.5 µl |
| Primer pool 1 or 2 (10 µM) | 3 µl |
| Q5 High-Fidelity DNA Polymerase | 0.25 µl |
| Nuclease-Free Water | 11.25 µl |
| Template DNA | 5 µl |

| **Step** | **Temperature** | **Time** |
|---|---|---|
| 1 cycle | 98°C | 30 seconds |
| 35 cycles | 98°C | 10 seconds |
| | 64°C | 3 minutes |
| 1 cycle | 72°C | 2 minutes |

**S3 Appendix.** Reaction setup and conditions for pool ITS2 (amplifying nuclear ITS2 gene)

| **Component** | **Amount for 25 µl reaction** |
|---|---|
| 5X Q5 Reaction Buffer | 5 µl |
| 10 mM dNTPs | 0.5 µl |
| Primer Ae_alb_ITS2_F (10 µM) | 1.25 µl |
| Primer Ae_alb_ITS2_R (10 µM) | 1.25 µl |
| Q5 High-Fidelity DNA Polymerase | 0.25 µl |
| Nuclease-Free Water | 11.75 µl |
| Template DNA | 5 µl |

| **Step** | **Temperature** | **Time** |
|---|---|---|
| 1 cycle | 98°C | 30 seconds |
| 35 cycles | 98°C | 10 seconds |
| | 64°C | 30 seconds |
| | 72°C | 1 minute |
| 1 cycle | 72°C | 2 minutes |
